# Rethinking Kalman Filters for Motor Brain-Machine Interface: The Fundamental Limitations and A Perspective Shift

**DOI:** 10.1101/2025.07.10.664247

**Authors:** Guanting Liu, Ying Yan, Jun Cai, Edmond Qi Wu, Shencun Fang, Adrian David Cheok, Aiguo Song

## Abstract

The Kalman filter has been introduced to the motor brain-machine interface (BMI) field for over 20 years and remains one of the most widely used models due to its simple and intuitive nature. However, this paper demonstrates that the application of Kalman filters in the BMI field violates the model’s own assumptions and numerous neuroscience principles, resulting in six critical limitations: (1) the observation matrix mapping from kinematics to neural activity causes individual neurons to be modeled independently, ignoring neural population activity; aligned kinematic and neural activity sequences fail to capture preparatory information; (3) the presence of behaviorally irrelevant neural activity causes observation weights to be excessively underestimated; (4) noise in the observation matrix renders posterior covariance estimates biased; (5) Kalman gain computation in observation space leads to repeated noise accumulation and (6) computational inefficiency. All these problems can be resolved through a simple perspective shift: separating the decoder from the Kalman filter and treating its predicted kinematics as the Kalman filter’s observation rather than neural activity. Experiments conducted on CRCNS-recorded monkey dorsal premotor area and primary motor signals performing computer cursor control tasks validate the existence of the six limitations and their violations of neuroscience principles, while demonstrating the superiority of the improved Kalman filter across all evaluated metrics.

## Introduction

Motor brain-machine interface (BMI) systems establish communication channels between subjects’ neural activity and external neuroprostheses, thereby offering potential for restoring motor function in patients with paralysis, amputation, amyotrophic lateral sclerosis, and other motor impairments (1, 2). The typical BMI workflow begins with the implantation of multi-array electrodes (MAE) in the motor cortex, usually in the dorsal premotor area (PMd) and primary motor cortex (M1) of experimental subjects including humans, rhesus macaques, and rats. These electrodes capture intracortical recordings that contain action potentials from individual neurons. Concurrently, the system records kinematic parameters from neuroprosthetic devices under manual subject control, such as computer cursors operated via joystick manipulation. Mathematical models are then developed to characterize the parametric relationship between neural activity and kinematics, establishing a predictive mapping from neural signals to intended movements (3, 4). This initial parameter learning phase with actual label is termed Manual Control (MC). Subsequently, subjects transition to direct neural control, where brain signals alone drive the neuroprosthetic device without manual input—a phase designated as Brain Control (BC), as shown in the figure 1. While some alternative experimental paradigms enable direct progression to BC for subjects incapable of performing MC (5, 6), the fundamental requirement remains the same: learning the complex parametric relationship between neural activity and intended kinematics.

**Fig. 1.**
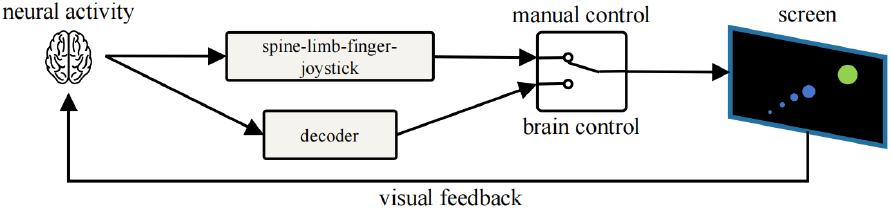
The workflow of a typical BMI system.

The Kalman filter is a powerful recursive state estimation algorithm whose core strength lies in optimally combining predictions based on system dynamics with noisy measurement to estimate the true state of a system. Its applications span an exceptionally broad range of domains, from GPS/INS navigation systems where it enables computationally efficient integration of satellite positioning and inertial measurement data (7), to robotic vision where it addresses uncertainties in localization, tracking, motion control (8), and medical text classification n the construction of clinical decision support systems (9). This versatility stems from the algorithm’s fundamental ability to handle uncertainty and noise in dynamic systems, making it an indispensable tool across diverse fields requiring real-time state estimation and prediction capabilities.

The inherent capability of Kalman filters to handle dynamic systems and noisy measurement data renders them exceptionally well-suited for BMI applications. In BMI system modeling, neural activity serves as the observation while kinematics represent the state, typically encompassing position, velocity, and acceleration variables. Prior to operational deployment, the system requires training in MC mode to establish critical parameters: the observation matrix, which typically employs linear regression to map kinematics to neural activity; the state transition matrix, which governs kinematic evolution according to physical principles; and both observation and process noise covariance matrices derived from empirical samples. Once these parameters are established, the system enables continuous motion control through recursive state estimation (10, 11).

Since its introduction to the BMI field by (12), the Kalman filter has become the one and only cornerstone algorithm in motor control, with applications rapidly expanding to enable patients with tetraplegia to control computer cursors, robotic arms, and other specialized devices (13– 17). Beyond motor applications, similar principles have been adapted to predict chronic pain states (18), while Kao et al. (19) demonstrated that neural population responses exhibit intrinsic dynamics that can be effectively captured through state-space models. When implementing Kalman filters for motor BMI systems, Closed-Loop Decoder Adaptation (CLDA) designs, where the brain acts as a controller selecting neural commands based on feedback of the current prosthetic state, represent a critical innovation for addressing neural signal variability. Orsborn et al. (20) demonstrated how continuous parameter recalibration during online control maintains performance regardless of initial decoder conditions, while Willett et al. (21) revealed important insights about user feedback control policies during operation. Mulliken et al. (22) enhanced state estimation by incorporating target information, showing how contextual knowledge can improve parameter learning. Among CLDA implementations, the ReFIT (Recalibrated Feedback Intention-Trained) Kalman filter (23) emerged as a landmark approach through fitting the decoder against estimates of intended velocity rather than actual movements, and explicitly modeling velocity as the primary intention while accounting for cursor position’s effect on neural activity. This framework became the standard against which new BMI approaches were measured, with applications to finger movement control (24), brain-computer typing interfaces (25), and investigations into neural variability (26, 27). Nason et al. (28) combined ReFIT with spiking-band power analysis to reduce power consumption, while (29) revealed how “covert rehearsal” can facilitate subsequent motor performance. Researchers also explored nonlinear Kalman filter variants to better capture neural-movement relationships. Li et al. (30) implemented an Unscented Kalman Filter with quadratic neural tuning, and Shoham et al. (31), Brockwell et al. (32) developed particle filtering approaches for recursive Bayesian decoding of non-Gaussian neural signals. Recent advances have further evolved Kalman filter implementations through integration with complementary approaches. Brandman et al. (33) proposed steady-state Kalman filter for rapid calibration, Abbaspourazad et al. (34) separated their model into jointly trained manifold and dynamic latent factors to capture non-linearity while maintaining tractable dynamics, and Haghi et al. (35) enhanced BMI control by integrating convolutional neural network feature extraction with conventional Kalman filtering.

However, is such a widely used model necessarily reasonable? This paper will demonstrate in detail in Section III Methodology that when transferring the Kalman filter, originally a control theory method, to motor BMI decoding, its assumptions are violated and its design contains flaws, including:

1. **Limitation 1:** The emission probability model maps from kinematics to neural activity, causing individual neurons to be modeled independently while ignoring neural population activity;
2. **Limitation 2:** Since the observation matrix is a mapping with compressed information that aligns kinematic sequences with neural activity sequences, the hidden Markov property cannot indirectly transmit historical information such as preparatory activity;
3. **Limitation 3:** The Kalman filter’s state estimation represents a mixture between process noise and observation noise, but the presence of behaviorally irrelevant neural activity causes observation noise to be overestimated, leading the Kalman filter to place insufficient trust in observations;
4. **Limitation 4:** Since the observation matrix is obtained through parameter learning, it inevitably contains noise (different from observation noise). The presence of this noise means that the Kalman filter’s posterior state estimation may not be unbiased, and the posterior covariance estimation is necessarily biased;
5. **Limitation 5:** The Kalman filter’s estimation process involves two dimensional transformations, both involving the noisy observation matrix, causing noise to accumulate multiple times;
6. **Limitation 6:** The calculation of Kalman gain occurs in the observation dimension, which is typically very high, resulting in excessive computational burden.

Before that, Section II will first review the relevant neuroscience knowledge and summarize three principles as supporting evidence for the subsequent argument.

## Preliminary

Motor BMIs aim to decode neural activity for device control, relying fundamentally on understanding the spatiotemporal dynamics of cortical motor systems. Early studies focused on correlating single-neuron activity with movement parameters but revealed profound complexity—individual neurons exhibit multiphasic responses, inconsistent directional tuning across contexts, and patterns distinct from muscle activation profiles (36, 37). This heterogeneity implies that single neurons cannot unambiguously encode movement kinematics, necessitating analysis of population-level dynamics where coordinated neural interactions generate reproducible low-dimensional trajectories (11, 38). Modern recording technologies, ranging from Utah arrays (hundreds of electrodes) to high-density systems like Neuralink’s 3,072-channel platform (39, 40) and the Argo system’s 65,536 channels (41), now enable granular tracking of these population dynamics across cortical regions.

Motor control involves distributed neural processing where preparatory activity in premotor cortex (PMd) begins hundreds of milliseconds before movement onset, establishing initial states that evolve into execution patterns in primary motor cortex (M1) (42, 43). These cortical commands undergo corticospinal transmission delays (80–150 ms) before reaching muscles (44, 45). Recorded neural signals integrate preparatory inputs, ongoing execution commands, and feedback adjustments based on sensory reafference (37, 46). This creates substantial temporal dependencies where kinematics reflect the integration of past and current neural activity.

Decoding this temporally extended activity faces two challenges. First, the neural-kinematic relationship is inherently statistical rather than causal. Parametric BMI models learn associations through training but remain susceptible to noise and non-determinism. For instance, erroneous prosthetic movements may still align with the subject’s internal predictions despite kinematic mismatches (46, 47). Second, recorded population activity contains substantial behaviorally irrelevant components. Motor cortical activity represents a mixed signal system where only a portion directly corresponds to movement output, while much reflects internal computations supporting movement generation. These processes include assembling movement “basis sets”—selecting and combining motor elements from broader pattern libraries to construct required commands. Though essential for movement generation, such internal computations lack direct correspondence with final movement parameters, appearing as noise in decoding tasks. Additionally, motor cortex continuously receives inputs from distributed brain regions, many unrelated to the current motor task, including background activity from attention networks, cognitive control systems, and other sensorimotor circuits (37, 47). This information is defined as behaviorally irrelevant neural activity (48–50).

In conclusion, the above content can be summarized into three neuroscience principles:

1. **Principle 1:** Individual neurons lack movement-specific encoding, necessitating multi-channel recording to capture relationships within neural population activity;
2. **Principle 2:** Kinematic features arise from preparatory, action, and feedback-related neural states spanning hundreds of milliseconds, implying that kinematics at any given moment correlate with both preceding and concurrent neural activity;
3. **Principle 3:** Cortical recordings represent a mixed signal system where substantial portions of recorded neural activity consist of behaviorally irrelevant components.

These assumptions will serve as supporting evidence for the subsequent section.

## Methodology

In this section, we will derive 6 limitations by combining the 3 principles with 4 perspectives: Hidden Markov Model, Gaussian Multiplication, Recursive Least Squares Estimation, and Linear Space Transformation. We will then provide improvement methods.

### A. Hidden Markov Model/Bayesian Perspective of Kalman Filter

The Kalman Filter can be viewed as an extension of Hidden Markov Models. Given *T* ordered random variables, according to Bayes’ theorem, their joint distribution can be written as a product of conditional distributions:

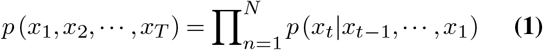

This product is often intractable in practice due to its enormous computational burden. If we directly introduce the Markov property *p* (*x*_*t*_ |*x*_*t−*1_, *· · ·, x*_1_) = *p* (*x*_*t*_ |*x*_*t−*1_), it would oversimplify the situation, ignoring the relationships between the current variable and earlier variables. Therefore, the idea of Hidden Markov Models is introducing hidden variables, allowing the variable at time *t* to connect with variables from previous time steps through the hidden variables. Defining the variables as *y* and hidden variables as *x*, we have the joint distribution of the Hidden Markov Model:

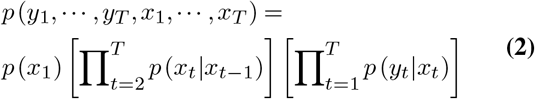

This simultaneously satisfies *x*_*t*+1_ ⊥⊥*x*_*t−*1_| *x*_*t*_. There-fore, the entire inference process is decomposed into three elements: the initial probability model *p* (*x*_1_), the transition probability model *p* (*x*_*t*_ |*x*_*t* −1_), and the emission probability model *p* (*y*_*t*_ | *x*_*t*_). In the Kalman Filter, the emission probability model corresponds to the observation equation, and the transition state model corresponds to the process equation:

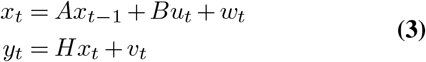

The control input *u*_*t*_ generally does not exist, so the process equation simplifies to *x*_*t*_ = *Ax*_*t−*1_ + *w*_*t*_. Both *w*_*t*_ and *v*_*t*_ are zero-mean Gaussian noise with covariances *Q* and *R*, respectively.

However, in the motor BMI field, new conditions are introduced. Unlike traditional Kalman filter applications, in motor BMI, the relationship between *y* and *x*, namely the observation matrix *H*, needs to be learned from multiple samples through parameter estimation beforehand. Typically, linear regression with least squares is used:

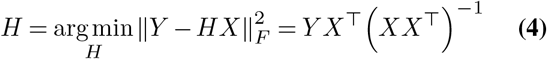

where *Y* ∈ℝ *q*×*N, X* ∈ℝ *p*×*N*. *p* is the dimension of the state vector, including prosthetic kinematics (such as position, velocity, acceleration). *q* is the dimension of the observation vector, generally the number of recorded neurons. *N* is the number of training set samples for parameter estimation.

This leads to **Limitation 1**. The design of the emission probability model (observation equation) makes the linear regression mapping direction from state vector to observation vector, which is equivalent to *q* independent linear regression equations, where the first one is:

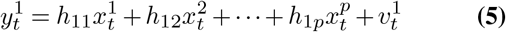

This means that each dimension of the observation vector (each neuron’s activity) is separately regressed by the state (kinematics), with neurons being independent of each other, which violates the **Principle 1**, that effective motor decoding should combine neural population activity. Even though this design uses multiple neurons, the relationships between neurons (such as linear combinations) are not modeled.

Next comes the **Limitation 2**. In the Hidden Markov Model framework, the state *x*_*t*_ at time *t* is indirectly connected to all previous observation vectors *y* through the transition probability model and emission probability model. This is unproblematic when the observation vector is a complete observation of the state vector system, where the observation vector *y*_*t*_ at time *t* contains all the information needed by the system.

However, this is unreasonable when the observation matrix *H* is learned through parameter estimation. According to the **Principle 2**, the motor state *x*_*t*_ at time *t* is related to the observation *y*_*t*_, *y*_*t−*1_… at time *t* and several previous time steps. Does the HMM framework “indirectly” capture this relationship? The answer is no. Because the observation matrix learned through aligned neural activity and kinematics sequence mapping involves information compression. The observation matrix at time *t* only models the relationship between *x*_*t*_ and *y*_*t*_; the information in *y*_*t*_ for *x*_*t*+1_, *x*_*t*+2_… after time *t* is not modeled, and similarly, at time *t −* 1, it models the relationship between *x*_*t−*1_ and *y*_*t−*1_. Therefore, even if the information from observation vector *y*_*t*_ is passed to *x*_*t*+1_ through hidden state transition (transition probability model), it is the information related to *x*_*t*_ in *y*_*t*_ (action activity), not the information between *y*_*t*_ and *x*_*t*+1_, *x*_*t*+2_… (preparatory activity).

Using *I*(*y*_*t*_ |*x*_*t*_) to represent information in *y*_*t*_ related to *x*_*t*_, as shown in the figure 2, black lines correspond to information successfully transmitted by HMM, showing that only all action activity is transmitted, while all preparatory activity (grey lines) is ignored. Ignoring these direct information pathways will cause the rich predictive information contained in neural activity to not be fully utilized, leading to degraded decoding performance.

**Fig. 2.**
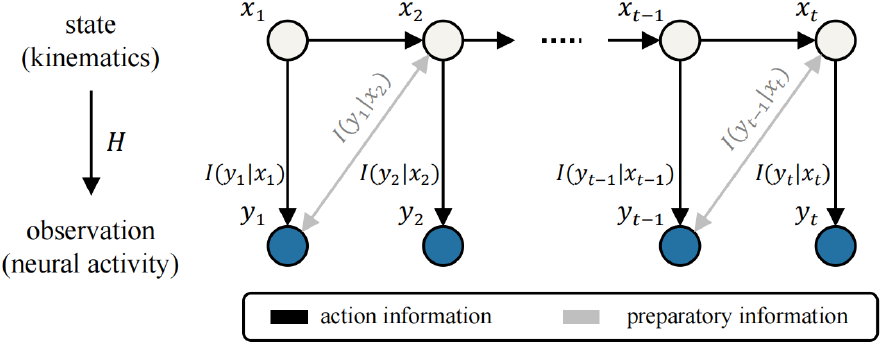
A typical HMM framework in brain-machine interface scenario.

So, can we achieve non-aligned learning by directly concatenating historical neural activity in each time step’s observation vector? The answer is no, and the solution to this problem will be provided after the introduction of the next section.

### B. Gaussian Multiplication Perspective of Kalman Filter

The Kalman Filter can also be understood from the perspective of Gaussian multiplication. It assumes that both the real observation and the state predicted by the transition equation (the prior state) follow a Gaussian distribution (also the observation decoded by prior state using observation matrix). The state (posterior state) finally output by the Kalman Filter combines the uncertainties of two Gaussian distributions, fusing them to obtain a Gaussian distribution with smaller uncertainty. For example, if two distributions have means and covariances of *µ*_1_, Σ_1_, *µ*_2_, Σ_2_, respectively, then the mean of the new distribution after Gaussian multiplication is:

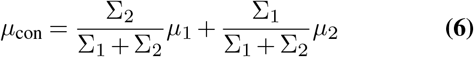

where the distribution with larger covariance corresponds to smaller weight.

Applying this to the Kalman Filter, first estimate the distributions of two noises using the training set:

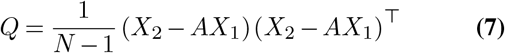

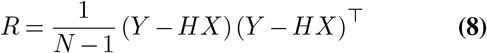

Then at time *t*, the Kalman Filter uses the state transition matrix to obtain the prior estimate of the state value at time t and its covariance:

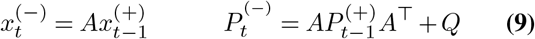

Since the state transition matrix *A* is linear, the output of this process is also a Gaussian distribution. Next, use the observation matrix to map it to the observation space to obtain the first distribution, the process distribution:

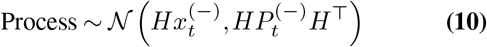

Meanwhile, the second distribution is the distribution corresponding to the observed value obtained at time t:

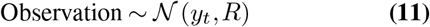

The Kalman Filter then fuses them:

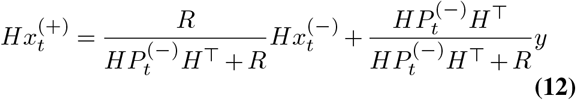

After simplification, we can obtain the classic iterative formula of the Kalman Filter:

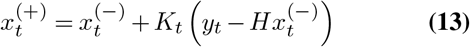

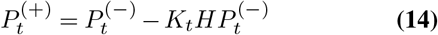

where:

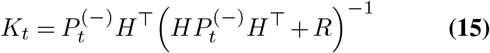

The Kalman gain serves as a tool for balancing the process distribution and observation distribution. If the observation noise *R* is low, then *K*_*t*_ approaches 1, and the posterior estimate trusts the latest observation *y*_*t*_ more, vice versa.

Consequently, **limitation 3** becomes apparent. According to **Principle 3**, when subjects perform motor control tasks, only a portion of their neural activity at this moment is related to motor control, while the remaining majority is irrelevant. Therefore, when the emission probability model *y*_*t*_ = *Hx*_*t*_ +*v*_*t*_ learns the target *y*_*t*_ through the controlled state *x*_*t*_, there must be a considerable portion that cannot be modeled. Even if the estimation of the observation matrix *H* is sufficiently accurate, achieving perfect modeling of the behaviorally relevant neural activity in the observed value *y*_*t*_, the remaining irrelevant activity portion will also cause the observation noise *v*_*t*_ and its covariance *R* to be high, making the Kalman Filter excessively distrust the observation distribution in the posterior estimation.

More formally, assume the observation vector *y*_*t*_ can be decomposed into two parts 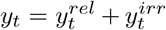, where 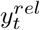 is the neural activity part related to motor control, and 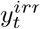 is irrelevant. Then ideally, the observation equation should only model the relevant part 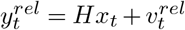, where 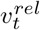 is the observation noise of the relevant part, with covariance *R*^*rel*^, which should typically be relatively low.

However, the actual observation equation can only uses the complete observation, where 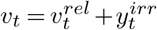, i.e., the observation noise actually includes 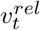 and the irrelevant neural activity 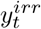. Therefore, the actual observation noise covariance is:

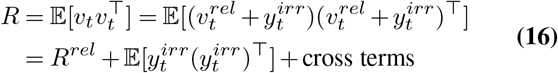

Assuming 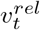 and 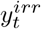 are uncorrelated, the cross terms are zero, then *R* = *R*^*rel*^ + *R*^*irr*^, where 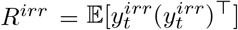 is the covariance matrix of irrelevant neural activity. Since 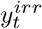 typically accounts for most of the neural activity, *R*^*irr*^ is often large, causing the overall observation noise covariance *R* to be high, resulting in *K*_*t*_ approaching zero. This means the Kalman Filter will greatly ignore observation information when updating the posterior estimate, over-relying on the prior estimate from the state transition equation 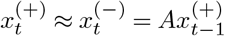:

In this situation, the Kalman Filter actually degenerates into a predictor that almost completely relies on the motion model, unable to effectively utilize the motor-related information contained in neural activity.

We now can answer the question mentioned in the previous part—”Can we achieve non-aligned learning by directly concatenating historical information in each time step’s observation vector?” From the Gaussian multiplication perspective in this part, we know that the observation contains a large amount of unlearnable parts (behaviorally irrelevant neural activity). If we expand the observation to include historical information, it would only further increase the variance (unlearnable parts), especially those observations more distant from current time *t*, which have lower correlation with the state at this moment and provide less information gain. Therefore, expanding the observation vector would actually lead to performance degradation. It might even directly cause problems like excessively large matrix condition numbers and ill-conditioned matrices during the observation matrix learning process.

### C. Recursive Least Squares Estimation Perspective of Kalman Filter

Kalman filtering can also be viewed as a recursive least squares estimation. When a new observation arrives at time *t*, the Kalman filter selects the Kalman gain that minimizes the posterior covariance to weight this new observation.

It can be proven that it is an unbiased estimate of the state, but this implicitly assumes that the observation matrix *H* is noise-free. Traditional control theory applications generally satisfy this condition. For example, in robot localization, the state vector consists of position, orientation, linear velocity, angular velocity, etc., while the observation vector consists of velocity measurements provided by wheel encoders, position measurements provided by GPS, etc. In this case, the observation matrix represents direct physical relationships between variables in the state vector and observation vector, so the observation matrix is noise-free. However, in motor BMI, the observation matrix *H* is estimated through training on multiple samples, which inevitably contains noise. Moreover, as demonstrated in the many limitations derived earlier, this noise will be very significant and may even have non-zero mean. Even if this observation matrix is learned well enough to have zero-mean noise, it will still make the posterior estimate of the covariance not an unbiased estimate. The following is the proof process:

Let the prior estimated state obtained at time *t* without measurement value *y*_*t*_ for correction be 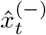, and the posterior estimated state obtained with measurement value *y*_*t*_ for correction be 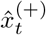. Let the observation matrix obtained from training set parameter estimation be 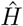. Let the theoretically completely noise-free observation matrix be 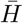, then 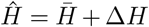. Meanwhile, for observation noise *v*_*t*_, it can also be decomposed into the noise of the observation value itself *v*_*ob,t*_ and the noise part that the model failed to successfully model *v*_*m,t*_. *v*_*t*_ = *v*_*ob,t*_ + *v*_*m,t*_ therefore satisfies:

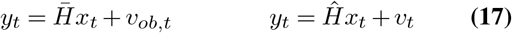

At this time, the error of linear recursive estimation is:

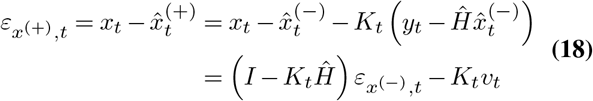

Then the covariance is:

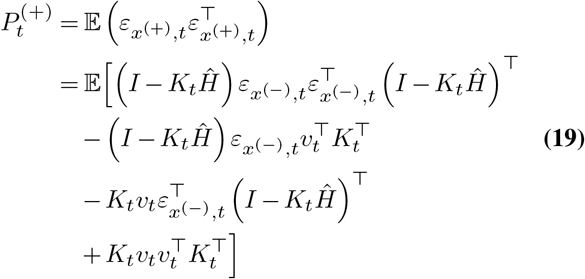

where the observation matrix *H* has fixed parameters during the inference period, so it is a constant, and:

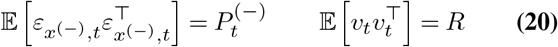

Therefore, the covariance can be simplified to:

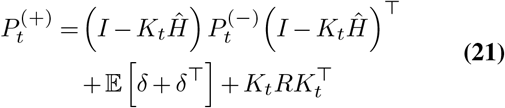

where:

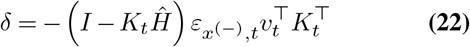

To minimize the error of the posterior estimate, the Kalman filter design seeks the optimal *K*_*t*_. We first ignore the variable *δ* and solve only for the remaining part:

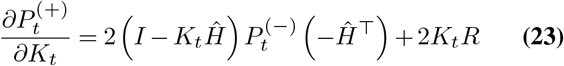

By setting 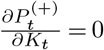, the solved *K*_*t*_ is optimal, yielding the basic formula of the typical Kalman Filter:

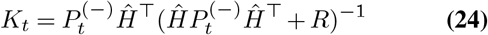

However, this is based on the case where *δ* does not exist. In actuality:

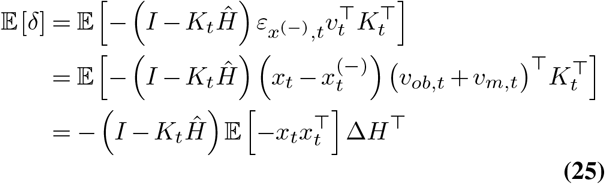

where *v*_*ob,t*_ has zero expectation and is independent of all other variables. *v*_*m,t*_ has zero expectation and is independent of 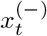 since it is the prior at time *t* that has not yer incorporated the observation:

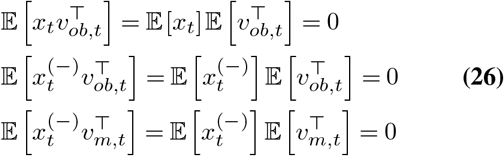

Obviously, 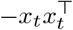 is positive semidefinite but not obtainable, so *δ* is ignored in typical Kalman filter in motor BMI, making it no longer an unbiased estimate.

The problem can be directly seen from the recursive formula of the posterior state from the perspective of recursive least squares estimation:

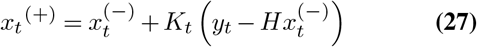

In this recursive formula, ideally, when multiplied by the noise-free *H* and compared with the new observation value *y*_*t*_ obtained at this moment, they match perfectly, so the correction term 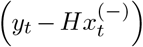 is 0, and the posterior estimate is exactly the same as the prior estimate. But if 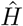 has noise, then even if the prior estimate is perfect, the correction term is not 0, leading to incorrect estimation of the posterior state. This is the **Limitation 4**.

### D. Linear Space Transformation Perspective of Kalman Filter

It can be seen that the state estimation of the Kalman filter undergoes two dimensional transformations. The first transformation maps the prior state to the observation space to obtain the “predicted” observation value of the prior state, and the second transformation maps this fusion value back to the state space as the posterior state estimate after fusing the “predicted” observation value of the prior state with the true observation value. The Kalman gain *K* serves as the mapping from observation space to state space. The dimension of the state vector *x* is ℝ*p*, and the dimension of the observation vector *y* is ℝ*q*. Generally, *q* is much larger than *p*, making it a tall matrix, and if the training of the observation matrix 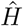 is ideal, it is full column rank. We can transform the iterative formula of states in the classic Kalman filter into a Gaussian multiplication perspective:

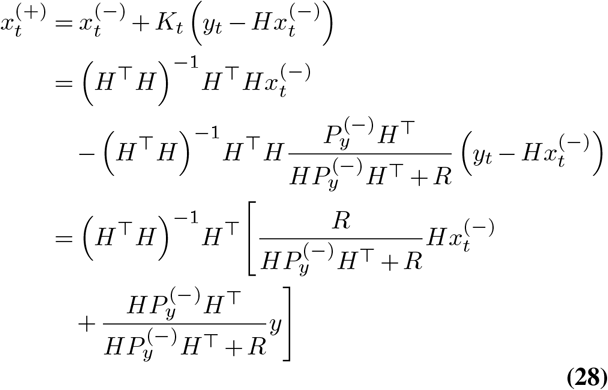

Since *H* is full column rank, *H*^*⊤*^*H* ^*−*1^*H*^*⊤*^*H* = *I*. Therefore, this equals the right side of equation (12) multiplied by the left pseudoinverse of *H, H*^*†*^ = *H*^*⊤*^*H* ^*−*1^*H*^*⊤*^. This is the least squares solution for *x* when *y, H* are known and *H* is not invertible, which is an approximate solution.

Since in motor BMI, the observation matrix 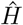 contains noise, errors are introduced when estimating the observation prediction value, and then when mapping this observation prediction value to the estimated state 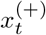 using the left pseudoinverse 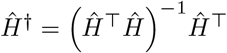, errors are introduced again. Therefore, this design is unreasonable in this scenario. This is the **Limitation 5**.

**Limitation 6** lies in the excessively high computational complexity. The Kalman gain is the step with the highest computational complexity in the iterative process, it is performed in the observation dimension. When performing single-step state estimation through standard iterative formulas (13), (14), and (15), the computational complexities are *O* (*pq*), *O* (*p*^2^*q*), and *O* (*q*^3^), respectively, which is inefficient when the number of neurons is very high.

If electrodes with thousands of channels like those mentioned earlier are used, the enormous computational cost would make real-time inference completely infeasible.

### E. Improvement Methods

#### E.1. Is the observation really neural activity?

Before proposing a solution, we must first reexamine the entire BMI system. This system includes MC (manual control, corresponding to the training phase) and BC (brain control, corresponding to the testing phase). Both convert neural activity into kinematics. The decoder in BC is a parametric model whose parameters need to be estimated from multiple samples in the training set by minimizing the error with the entire pathway in the MC phase. Although from a neuroscience perspective, the neural activity in the MC and BC phases differs, overall, the ideal state of BC is to “mimic” MC. If Kalman filter serves as this decoder, with neural activity as input and kinematics as output, then neural activity would naturally be the observation in Kalman filtering.

But is this really the case? The particularity of the BMI field lies in the fact that the relationship between neural activity and kinematics lacks empirical formulas and must be obtained through supervised learning-in this case, the observation matrix. This differs from traditional Kalman filter, which does not involve supervised learning and therefore lacks the step of “mimicking” the MC phase loop. In reality, this ‘mimic’ of MC is achieved entirely through the observation matrix, which BMI researchers have implemented as a linear regression matrix in their adaptation of Kalman filter. From this perspective, this linear regression matrix is the actual decoder, not Kalman filter. Kalman filter is positioned after this decoder, and its inference phase is separated from the linear regression matrix’s inference phase (corresponding to the fixed observation matrix parameters when using Kalman filtering).

Furthermore, we can reasonably argue that the truly appropriate observation for Kalman filter in the BMI field should be the output of the decoder (linear regression matrix)—that is, the predicted kinematics. In this case, both the state and observation of Kalman filtering exist in the kinematics space. While this may seem counterintuitive, I will demonstrate why it is more reasonable and how it can address the 6 limitations while conforming to the 3 principles.

#### E.2. Improved Kalman Filter

According to the above discussion, the linear regression matrix can now be separated independently, and the mapping method is from *y* to *x*:

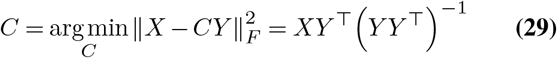

Making 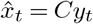. Meanwhile, the calculation of observation noise covariance is:

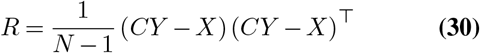

In this way, the observation value in this system is actually the state vector 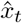 estimated through neural activity *y*_*t*_ at time t. At this point, the observation equation becomes:

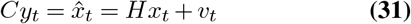

Then, the observation vector and state vector are unified. The observation matrix *H* becomes an identity matrix:

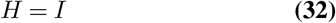

Based on this, we can rewrite the standard iterative formulas (13), (14), and (15):

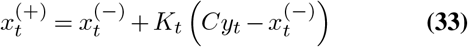

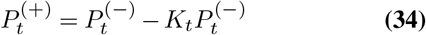

where:

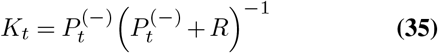

Next, we can gradually analyze why this simple design can solve the 6 limitations mentioned above. First, in the modified framework, the step linking neural activity with kinematics is placed outside the Kalman filter, and the direction has changed, where the estimate of the first kinematics component is::

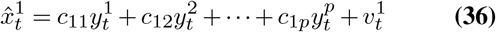

This solves **Limitation 1**. In this linear regression, each dimension of the state vector estimated from observations is a linear combination of all neuronal activities. Therefore, relationships between different neurons are considered, using neural population activity to estimate the state, rather than separately regressing each neuron’s activity from the state vector as in the previous design.

Second, regarding **Limitation 2**. As mentioned earlier, in the typical Kalman filter design, the state vector *x*_*t*_ at time *t* is directly connected to the observation vector *y*_*t*_ at time *t*, but indirectly connected to observation vectors before time *t*, and this “indirectness” cannot be achieved when the observation matrix is a linear regression matrix between the state vector *x*_*t*_ and the observation vector *y*_*t*_. In the improved framework, since the step *c* linking neural activity with state vectors is placed outside the Kalman filter, the observation vector within the Kalman Filter system is the state vector 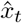 predicted by that step. Therefore, we can completely introduce observation values before time *t* when predicting 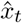 without modifying the Bayesian inference framework (Hidden Markov Model) of the Kalman filter:

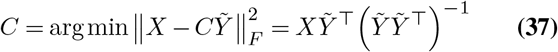

Making 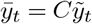, where:

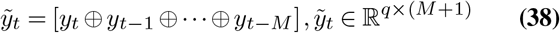

⊕ represents the vector concatenation operation. Therefore, the prediction of 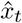 uses neural activity from *y*_*t*_ and its previous *M* time steps, directly linking the kinematics at time *t* with neural activity at time *t* and several previous time steps, as shown in Figure 3. Meanwhile, the observation covariance is:

**Fig. 3.**
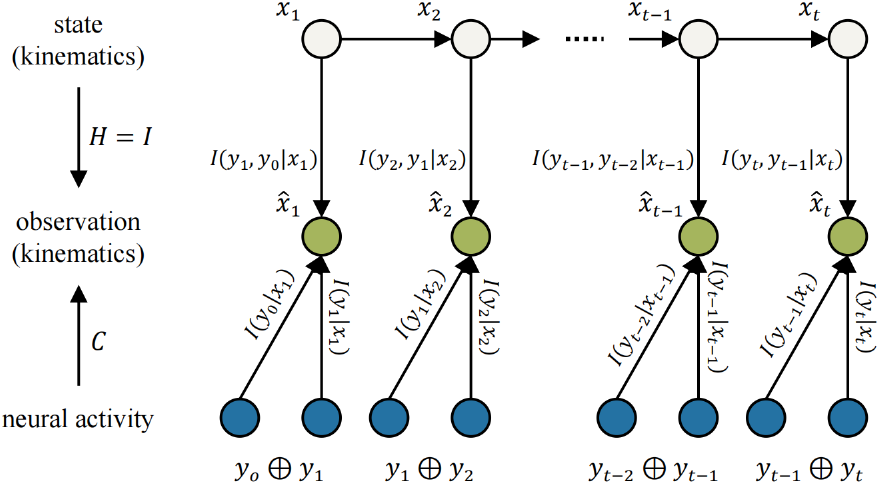
Framework of the improved Kalman filter. The HMM component remains unchanged, with the emission probability model still mapping from state to observation. However, the observation is now predicted kinematics rather than neural activity.

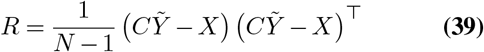

**Limitation 3** is also solved simultaneously. Because the mapping direction of linear regression is now from neural activity to kinematics, the parts in neural activity unrelated to kinematics are automatically compressed during this process, which will not cause unestimable components in the regression target.

Meanwhile, this approach is also more reasonable from the perspective of Gaussian multiplication. Kalman filtering is the fusion of process distribution and observation distribution, but in the typical Kalman filter design, as described in equations 10 and 11, the observation distribution is the distribution of true observed values, while the process distribution is the result of the prior state from the process equation being mapped to the observation (neural activity) space through linear regression. This means it conflates errors from both the process and observation components, which is unreasonable.

However, in the improved Kalman filter design, the observation matrix is an identity matrix *I*, completely noiseless. Therefore, the error after mapping to the observation (kinematics) space contains only errors originating from the process equation, while the observation distribution also contains only errors from the observation equation. Thus, fusing these two distributions is reasonable:

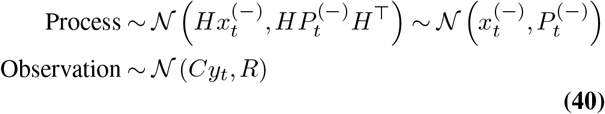

The solution to **Limitation 4** is more intuitive. In the modified Kalman Filter, the observation vector *H* = *I*, therefore completely noise-free, with noise only existing in the observation, making the estimates of posterior state 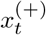 and posterior covariance 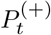 unbiased estimates.

Next is **Limitation 5**. The linear space transformation situation in the modified iterative formula is as follows:

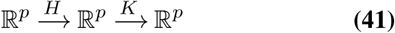

where *H* is noise-free, so these two mappings will not introduce additional noise. Meanwhile, for **Limitation 6**, since all operations of the Kalman Filter occur in the state dimension, the computational complexity is greatly reduced. At this time, dimensions *p* and *q* are equal, both being the dimension number of kinematics, which is a low-dimensional vector (5 in this paper). This iterative process is unrelated to the high-dimensional neural activity vector dimension (neurons), so no matter how many more neurons we add, it will not increase the iterative computational burden of the Kalman filter.

Thus, the six limitations can all be solved with this simple modification.

#### E.3. Extensibility of the Improved Kalman Filter

In the previous parts, we mentioned that since the observation of the Kalman filter is now the predicted kinematics from an external decoder, the decoder no longer needs to be typical linear regression, but can be replaced with other linear, nonlinear models or even neural networks, as long as its noise is Gaussian. To avoid excessive jumping, this paper demonstrates replacing the decoder with Partial Least Square Regression (PLSR), which is highly similar to linear regression (both ultimately requiring only a linear matrix for mapping) while offering superior interpretability. PLSR enables analysis of variable importance through regression coefficients and reveals the components of independent variables (neural activity) and dependent variables (kinematics) through interpretable components that emerge as the number of components increases.

The workflow of PLSR is shown in Algorithm 1. Given standardized data matrices *X* ∈ ℝ *p*×*N* and *Y* ∈ ℝ *q*×*N*, where *N* is the number of samples, *p* and *q* are the number of variables in *X* and *Y*, respectively. PLSR establishes a linear relationship *X* = *BY* between *X* and *Y* by extracting latent variables (components).

##### Algorithm 1

Partial Least Squares Regression

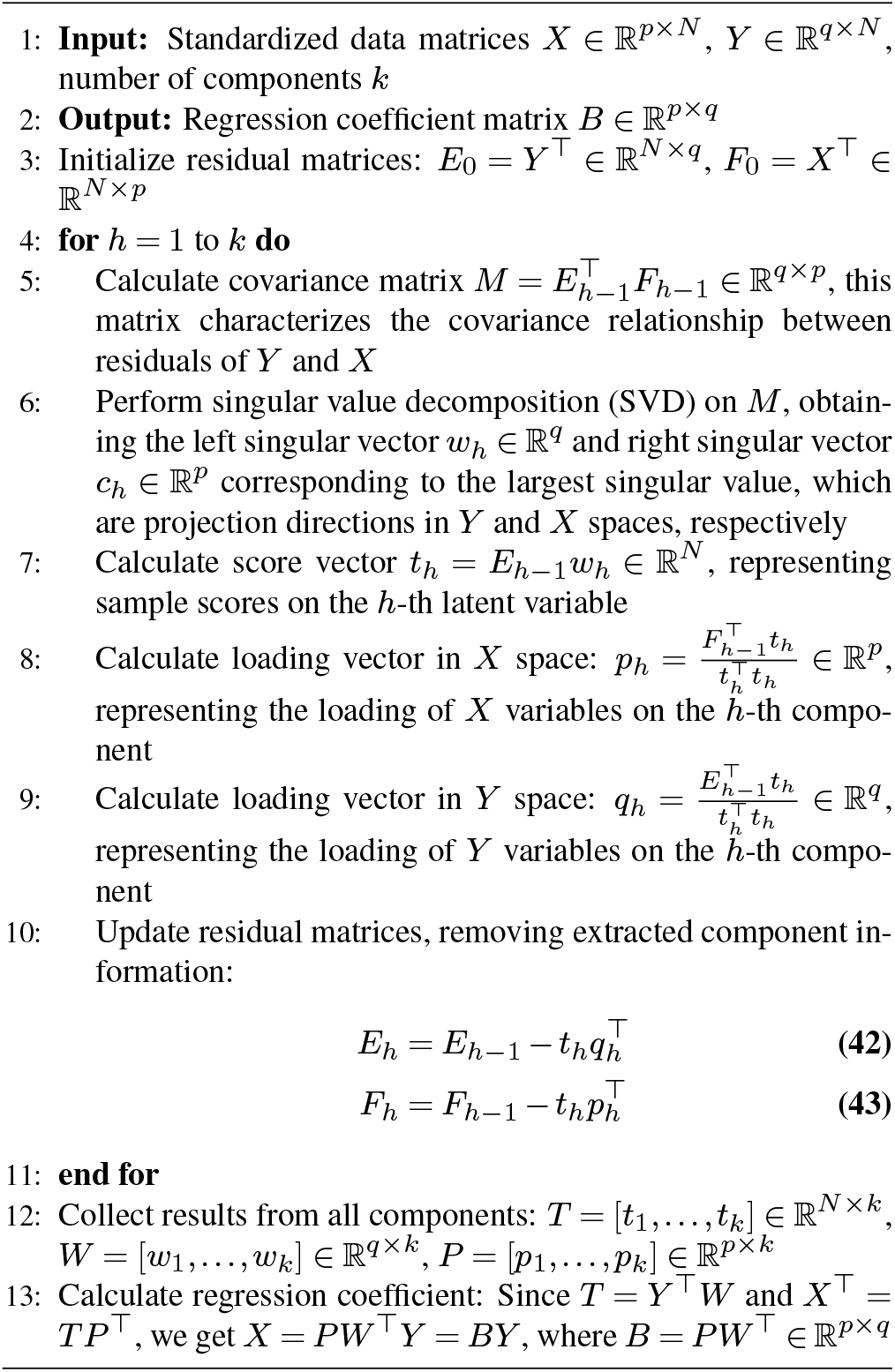

Each iteration of PLSR (with increasing components) explains partial correlated components between current *X* and *Y*, gradually converging when the remaining residuals no longer exhibit correlation. The final regression coefficient matrix *B* makes *X ≈ BY*. Its functionality will be utilized in the next section.

## Results

### A. Dataset and Experimental Setup

The dataset utilized in this research was made publicly available by Matthew G. Perich from Lee E. Miller’s laboratory at Northwestern University (51, 52), with all experimental procedures adhering to institutional animal care standards. Neural recordings were obtained from a macaque monkey fitted with two 100-electrode Utah arrays positioned in the primary motor cortex (M1) and premotor cortex (PMd). Manual spike sorting was performed using conventional methods.

During the experimental sessions, the monkey manipulated a cursor in a 20 cm × 20 cm workspace, executing reaching movements toward pseudo-random targets positioned within a 5-15 cm radius. Target locations appeared at arbitrary positions relative to the starting point, diverging from standard center-out paradigms. New targets emerged 200 ms following arrival at the current target, after a 100 ms holding period, creating smooth movements with variable distances. Neural spike trains were characterized as binned spike counts using non-overlapping 50ms time windows from 161 neurons across M1 and PMd regions. Cursor kinematics were sampled at 10 ms intervals, subsequently downsampled to 50ms for positional data, with velocities recalculated from these positions. The model’s trajectory output was reconstructed using predicted velocities rather than directly employing predicted positions, producing smoother movement paths.

The dataset encompasses recordings from two subjects (MM and MT), though only MM’s data was utilized since MT’s sessions contained exclusively PMd recordings lacking corresponding M1 data and demonstrated inconsistent neuronal correspondence between sessions. Initially, individual trials encompassed 1-4 consecutive reaching movements. To mitigate cumulative errors from disproportionately impacting extended trials, we partitioned each trial, treating individual reaches as distinct trials. This approach yielded 496 trials with durations spanning 750ms to 1680ms.

### B. Importance of Neural Activity from Different Sources

As described in the previous section, we adopt PLSR to replace linear regression because PLSR provides excellent interpretability for analyzing the neuroscience principles mentioned in the Related Works section. It can be used to analyze the importance of different neural activities, including contributions from different neurons, brain regions, and time points, as well as to analyze behaviorally irrelevant/relevant neural activity components.

While PLSR obtains a mapping from kinematics to neural activity similar to linear regression, it can also reflect variable importance through PLS-VIP (Variable Importance in Projection) scores and PLS-BETA scores (53).

The VIP score for the *j*-th variable is defined as:

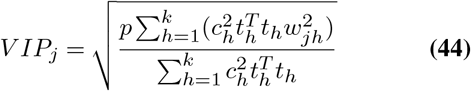

where *p* represents the total number of variables participating in the analysis; *k* represents the final total number of iterations performed; *w*_*jh*_ represents the *j*-th loading coefficient for the *h*-th dimension; 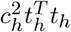 represents the degree of interpretation of the *h*-th component result *X*_*h*_ for *Y*_*h*_ (Note: *X*_*h*_ is the component result of independent variable *X* for the *h*th iteration, *Y*_*h*_ is the part of dependent variable *Y* interpreted in the *h*-th iteration).

The BETA score represents the regression coefficients of each variable in the final PLS regression model. Unlike ordinary least squares regression, the regression coefficients in PLS reflect the degree of correlation between variables after considering the covariance structure. Variables with larger absolute BETA values contribute more to the prediction (whether positive or negative correlation), while variables with smaller absolute BETA values contribute less. PLS-BETA provides a more stable coefficient estimation in multivariate analysis and can be used in conjunction with VIP scores to evaluate variable importance from different perspectives.

Figure 4a and 4b show the VIP scores and BETA scores with bin size set to 5, meaning that the kinematics at each time step are represented by a vector concatenating the binned spike counts from the current moment and the previous 4 moments (5 bins total) as independent variables for PLSR. Neural activities from two regions (PMd and M1) across 5 bins are grouped, and the VIP score distribution for each group is visualized. As shown in the figure, neural activities from PMd and M1 exhibit approximately equal values at each time step. The VIP and BETA scores across all time steps demonstrate information gain for kinematics, confirming the existence of preparatory neural activity. The current time step’s neural activity shows the highest VIP score, contributing most to kinematics regression. VIP scores decrease as time steps become earlier, and this trend is similarly reflected in the absolute values of BETA scores.

**Fig. 4.**
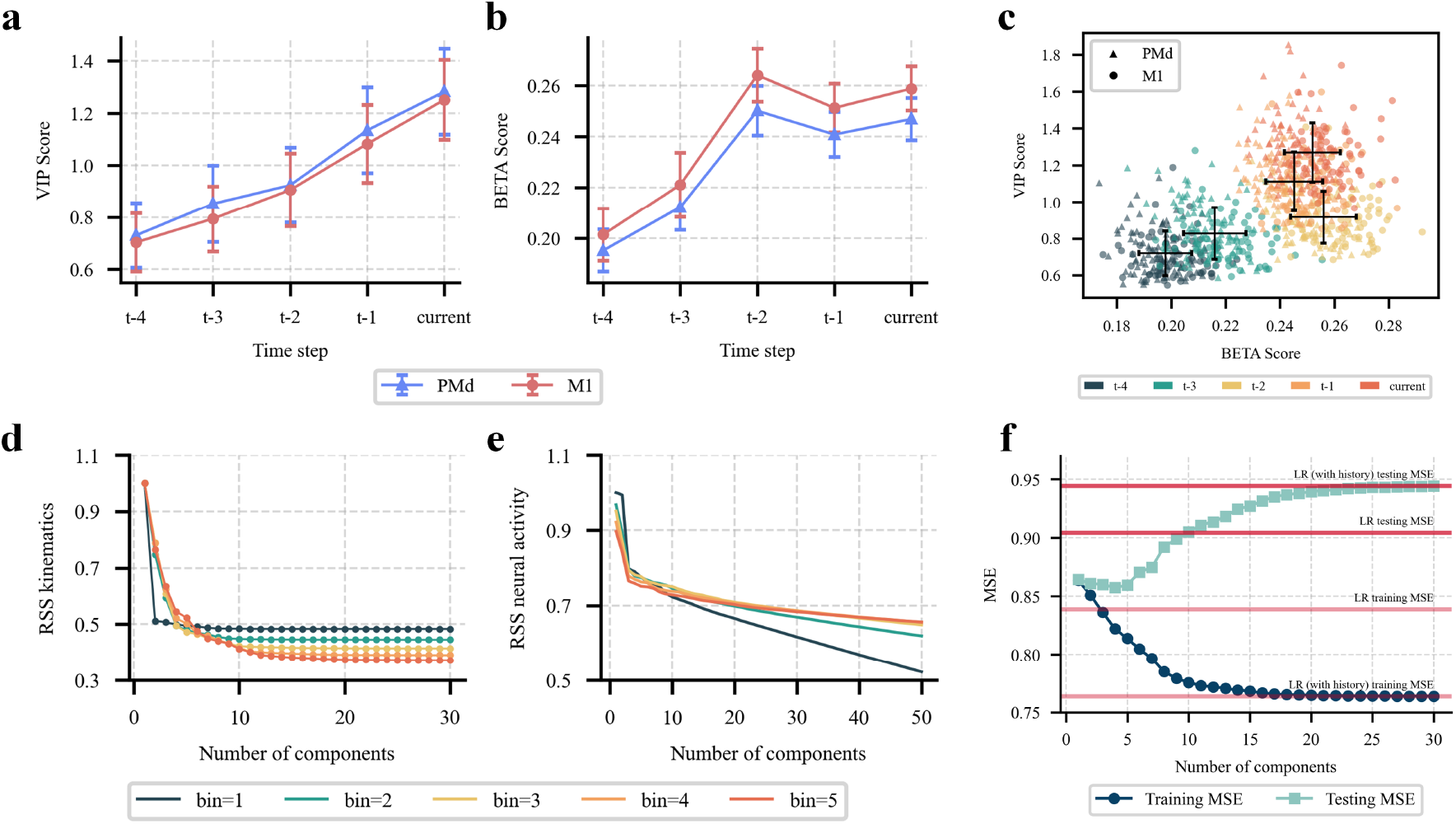
Analysis of neural activity importance and behavioral relevance using PLSR. **a**. VIP scores for neural activities from PMd and M1 regions across 5 time bins. **b**. BETA scores for neural activities from PMd and M1 regions across 5 time bins. **c**. Relationship between BETA and VIP scores across different time steps. **d**. Residual sum of squares (RSS) trends for kinematics during PLSR iterative process with different bin sizes. **e**. RSS trends for neural activity during PLSR iterative process with different bin sizes. **f**. Training and test MSE comparison between linear regression (LR) and PLSR with/without history.

This validates our answer to the question posed in the previous section: “Can we achieve non-aligned learning by directly concatenating historical information in each time step’s observation vector?”. Historical information indeed provides information gain for kinematics, but earlier neural activity has lower signal-to-noise ratios (lower score). Therefore, when regressing neural activity from kinematics, this approach would only decrease performance.

Additionally, the BETA score reveals a specific pattern: it peaks at time step t-2 (100 ms before current) rather than continuously decreasing with earlier time steps. This time point corresponds precisely to the preparatory neural activity period. Figure 4c plots BETA scores as the x-axis and VIP scores as the y-axis, showing that excluding the t-2 time point, VIP and BETA exhibit a linear relationship, while t-2 represents an exception.

### C. PLSR Iterative Process Reflects Changes in Behaviorally Irrelevant/Relevant Neural Activity

We can reflect the irrelevant components in kinematics and neural activity through the changes in residual sum of squares (RSS) during PLSR’s iterative process (increasing components), as the correlated parts extracted in earlier iterations are more substantial. Figure 4d and 4e show the declining trends of residual variance in kinematics and neural activity as the number of iterations (components) increases, using bin sizes from 1 to 5.

For kinematics trends, longer bins show slower decline rates, but all essentially stop declining when the number of components almost reaches 15, indicating that no neural activity components remain that can contribute further to kinematics regression—i.e., no more behaviorally relevant neural activity exists. However, the residual decline trend for neural activity differs from that of kinematics. Neural activity residuals continue to decrease with increasing component numbers, even when excess neural activity at 15 components no longer contributes, with more and more neural activity being continuously extracted into the latent space. From this perspective, the residual neural activity from the 15th component onwards can be regarded as behaviorally irrelevant neural activity, since their inclusion does not lead to further fitting of kinematics.

Figure 4f further demonstrates this phenomenon. The light red lines represent training set MSE for linear regression (LR) with and without history, respectively, while dark red lines represent the test set. Remarkably, although LR with history shows lower training set MSE than without history, it actually shows higher test set MSE, indicating overfitting. PLSR with history shows training and test set MSE in light cyan and dark cyan, respectively. The training set MSE continuously decreases with increasing components until it equals the training set MSE of LR with history. However, the test set MSE first slightly decreases then increases until it equals the test set MSE of LR with history. This indicates that selecting a relatively small number of components (e.g., 5) is sufficient, and excessively high PLSR component numbers will lead to overfitting (equivalent to LR).

### D. Linear regression from kinematics to neural activity

Next, we validate the reasonableness of the observation matrix in the conventional Kalman filter design, which represents a mapping from kinematic to neural activity. In this approach, Gaussian multiplication is computed in the observation space, and the result is then mapped back to the state space through the left pseudo-inverse of the observation matrix. Figure 5a demonstrates this result. Matrix 1 (*H*) represents the linear regression from neural activity to kinematics, while Matrix 2 (pseudo-inverse *H*) represents the left pseudo-inverse of the linear regression from kinematics to neural activity (corresponding to the conventional Kalman filter). After training, we used the test set’s neural activity to predict kinematics through both matrices, and calculated the mean squared error (MSE) against the ground truth kinematics, yielding four distributions (corresponding to position and velocity in x and y axes, respectively). Note that there are no negative values in the loss data. The negative regions in the violin plots are due to the smoothing effect of kernel density estimation, which causes the density curves to extend into negative regions even when all data points are positive. It is evident that the MSE of kinematics regression using pseudo-inverse *H* is significantly higher than that using *H* (p < 0.0001), demonstrating the inefficiency of regressing neural activity from kinematics.

**Fig. 5.**
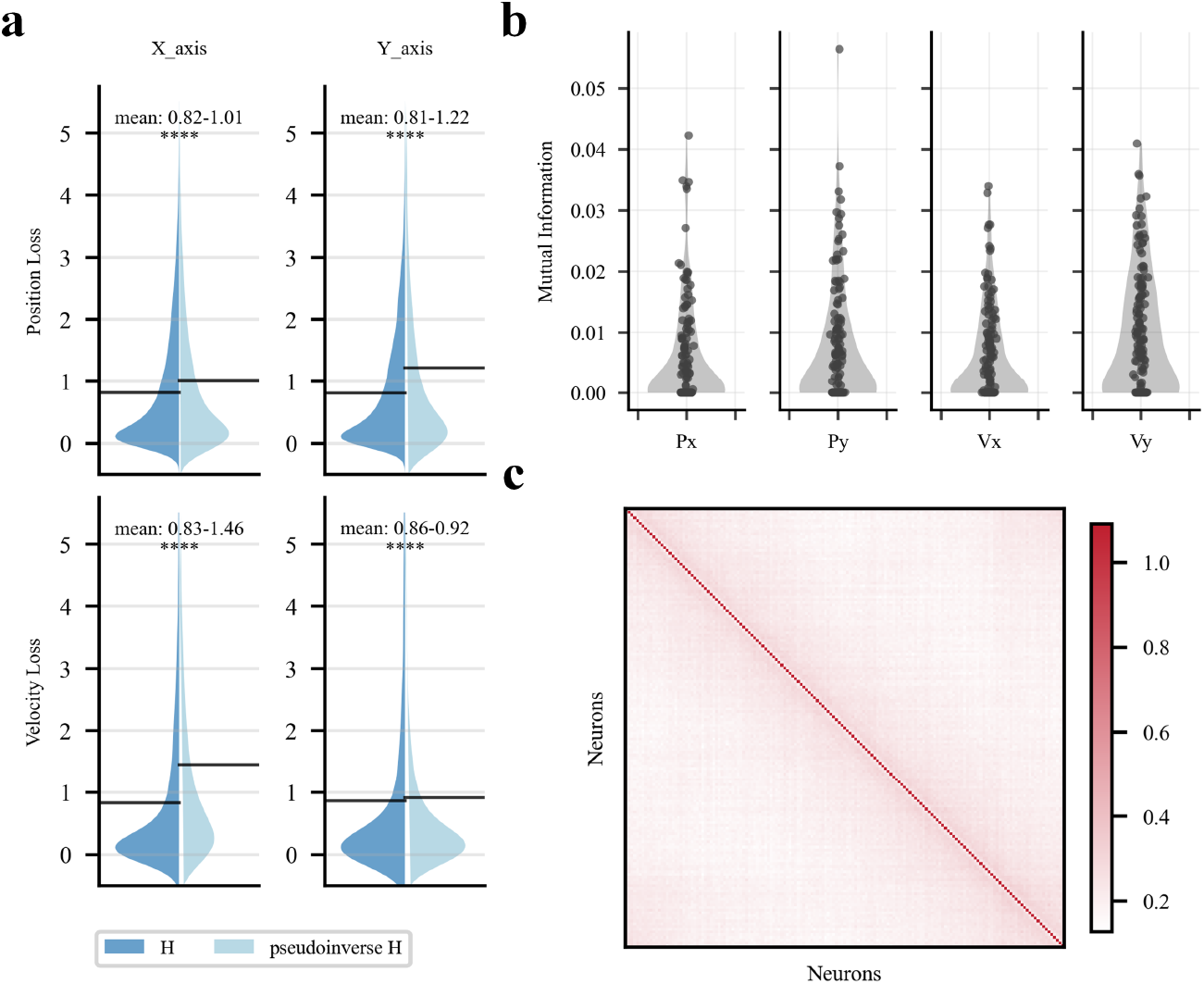
Validation of the observation matrix in conventional Kalman filter design. **a**. MSE distributions comparing kinematics prediction using Matrix 1 (*H*: neural activity to kinematics) versus Matrix 2 (pseudo-inverse *H*: kinematics to neural activity) across four kinematic dimensions. **b**. Mutual information between each kinematics component and individual neurons. **c**. Covariance matrix between predicted neural activity (from kinematics) and actual neural activity, corresponding to observation noise covariance in conventional Kalman filter.

This occurs because, as previously mentioned, neural activity contains substantial behaviorally irrelevant components that are unrelated to kinematics and cannot be learned through this linear regression. Consequently, using such activity as the regression target leads to poor performance. Figure 5b illustrates this result by showing the mutual information between each component of kinematics and all individual neurons. Most mutual information values are extremely low, indicating that a large portion of neural activity consists of irrelevant information. Further results are presented in Figure 5c, which shows the covariance matrix between the predicted neural activity derived from kinematics and the actual neural activity. This corresponds to the observation noise covariance in the conventional Kalman filter. Notably, the diagonal elements are nearly equal to 1, indicating that regressing neural activity from kinematics is essentially equivalent to random guessing.

### E. Performance Comparison

We compared the performance of four decoding methods: the conventional Kalman filter (with observation matrix mapping from kinematics to neural activity), state-based Kalman filter (with neural activity-to-kinematics decoder independent of the Kalman filter), state-based Kalman filter with history (bin = 5, incorporating 200 ms of historical neural activity), and Kalman filter using PLSR as the decoder (also with bin = 5). Consistent with other studies, we used the velocity predicted by the Kalman filter to generate trajectories, as this produces smoother curves. When calculating the difference between predicted and actual velocities, we used mean absolute error (MAE), obtaining one MAE value per trial, resulting in 496 MAE values total.

Figure 6a shows the MAE distributions for the four models, with the conventional Kalman filter performing worst, clearly inferior to all other methods (all state-based approaches). The performance of the four decoding methods was evaluated using paired t-tests to compare MAE values across all trials. Since the same neural data was used for all methods within each trial, paired comparisons were conducted to account for trial-to-trial variability. Normality of the data was assessed using the Shapiro-Wilk test, which indicated non-normal distributions for all methods. However, given the large sample size (n = 496 trials), the t-test remained robust due to the central limit theorem. Multiple pairwise comparisons were performed between all four methods (6 comparisons total), with Bonferroni correction applied to control the family-wise error rate (corrected *α* = 0.05/6 = 0.0083). Effect sizes were quantified using Cohen’s d to assess the practical significance of observed differences. Additionally, the False Discovery Rate (FDR) correction using the Benjamini-Hochberg procedure was applied as a less conservative alternative to Bonferroni correction.

**Fig. 6.**
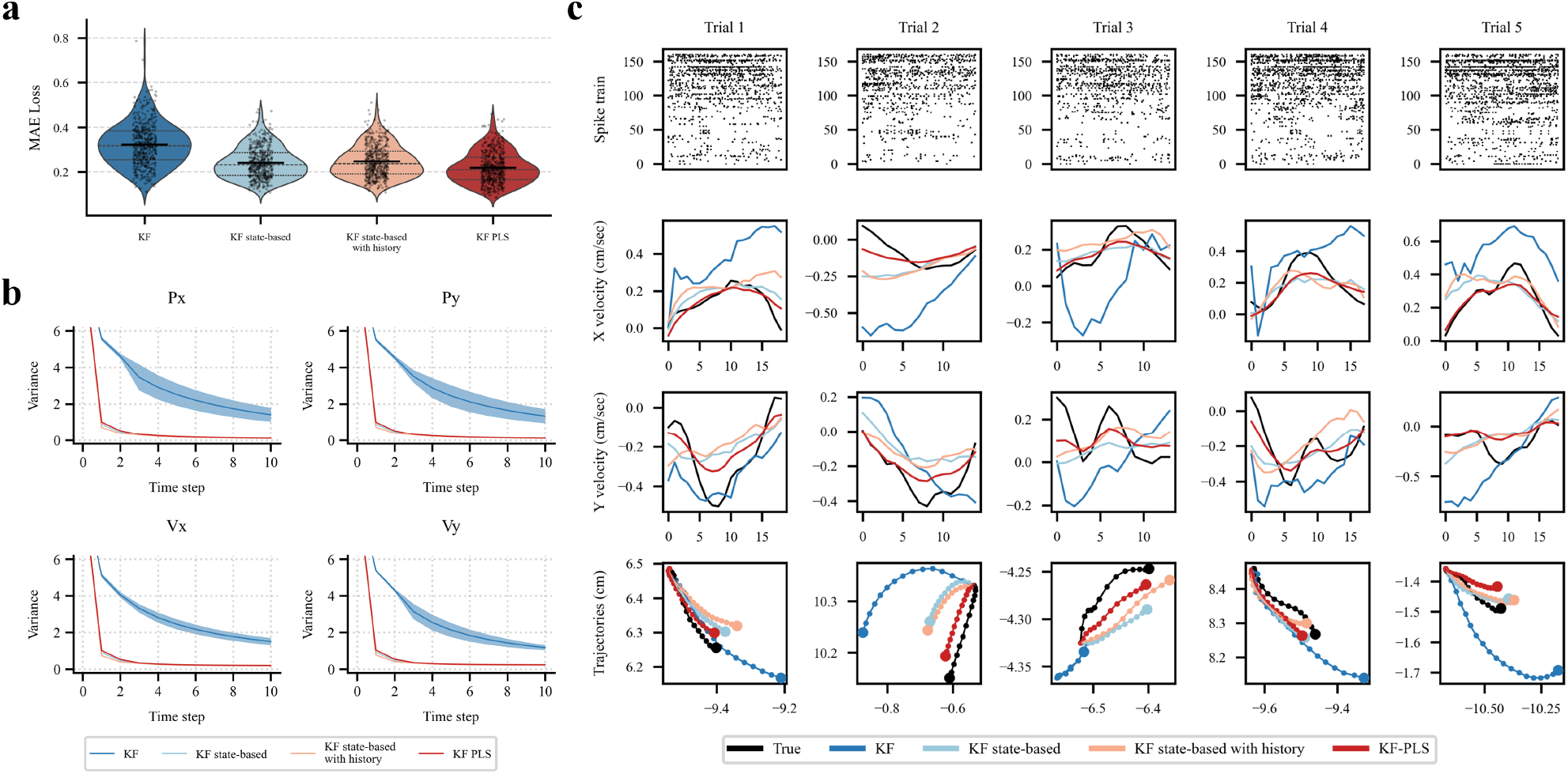
Performance comparison of four decoding methods. **a**. MAE distributions for conventional Kalman filter, state-based Kalman filter, state-based Kalman filter with history, and Kalman filter using PLSR as decoder. **b**. Changes in state variance (diagonal elements of state covariance matrix) for each kinematic variable during real-time estimation across the four models. **c**. Spike firing patterns for several trials showing PMd region (first 94 neurons) and M1 region (67 neurons), along with trial X and Y velocity curves and cursor trajectory in 2D space.

All pairwise comparisons revealed statistically significant differences (all p < 0.001, Bonferroni corrected), with large effect sizes (Cohen’s d ranging from 0.48 to 2.62). The KF-PLS method demonstrated superior performance with the lowest MAE (0.219 ± 0.071), followed by KF state-based (0.241 ± 0.071), KF state-based with history (0.247 ± 0.072), and conventional KF (0.323 ± 0.097). The KF-PLS method showed a 31% improvement in decoding accuracy compared to the conventional KF approach.

Additionally, Figure 6c displays the spike firing patterns for several trials, including the first 94 neurons corresponding to the PMd region and the subsequent 67 neurons corresponding to the M1 region. The firing rate in PMd is notably higher than in M1. The figure also shows the trial’s X and Y velocity curves as well as the cursor trajectory in the 2D computer space.

Furthermore, Figure 6b illustrates the changes in state variance (diagonal elements of the state covariance matrix) for each kinematic variable during real-time estimation across the four models. Except for the conventional Kalman filter, which converged extremely slowly and had not fully converged by the end of estimation, all state-based Kalman filters achieved rapid convergence. This demonstrates the phenomenon proven in previous sections: the conventional Kalman filter repeatedly accumulates noise during linear space transformation due to noise in the observation matrix.

Finally, as mentioned earlier, the Kalman gain computation in conventional Kalman filters occurs in the neural activity space, resulting in computational inefficiency. We tested the computational efficiency comparison between conventional and improved Kalman filters when inferring multiple trials. As the number of neurons increased from 30 to 150, the inference time of conventional Kalman filters increased almost linearly from 5 seconds to 32 seconds. In contrast, the improved Kalman filter was barely affected by the number of neurons, with computation time consistently fluctuating around 2.7 seconds.

## Discussion

Thus far, we have explored the 6 limitations in conventional Kalman filters and how they violate the 3 neuroscience principles. We have proposed that simply treating the predicted kinematics from linear regression as the observation in Kalman filter can resolve these limitations. Most importantly, traditional BMI applications of Kalman filters, in order to preserve their original properties, rely primarily on simple linear regression for the most crucial part of the entire system—decoding neural activity into kinematics. This approach fails to capture complex nonlinear and non-stationary relationships, greatly limiting the exploration and application of complex neural activity. The separation of Kalman filtering and decoder under our framework can break through this limitation, as the decoder can be any other model, requiring only that its output error follows a Gaussian distribution.

However, this raises another question. Since Kalman filter is now completely separated from the decoder, wouldn’t using only the decoder be sufficient to implement the entire process? Why is Kalman filtering still needed? This is actually a dilemma that this paper aims to highlight. Kalman filter has been introduced to the BMI field since 2003 and has remained the most commonly used model, but this paper argues that it is not an excellent model.

The main reason is that its model capacity (number of parameters) is too small. Kalman filter’s dependence on linearity, Gaussian distributions, and Markov properties restricts its development toward large-scale nonlinear models. As the brain is a highly nonlinear, dynamically changing system, large-scale data-driven methods should benefit it. The neuroscience and brain-machine interface fields have not yet experienced their large-scale data-driven AI revolution, which this paper considers to be the future research direction.

## Acknowledgments

This work was funded by the National Natural Science Foundation of China under Grant 52077105, in part by the Natural Science Foundation of Jiangsu Province under Grant BK20211285, in part by the Open Foundation of Key Laboratory of Technology and Equipment of Tianjin Urban Air Transportation System, under Grant TJKL-UAM-202301, and in part by Excellent Research and Innovation Teams in universities in Anhui Province, under grant 2023AH010021.

